# Macromolecular crowding by hyaluronan: implications for binding reactions in the pericellular matrix and at the cell surface

**DOI:** 10.1101/2024.01.03.573870

**Authors:** David G. Fernig

## Abstract

In common with many glycosaminoglycan binding growth factors, the interactions of fibroblast growth factor-2 (FGF-2) with heparin/heparan sulfate regulate its biological activity. Consequently, much effort has been expended in characterising growth factor-glycoaminoglycan interactions. Many experiments are performed <u>in vitro</u> in dilute solution. However, the cell surface is an extremely crowded environment. We have, therefore, examined the parameters of FGF-2 binding to a heparin-derived tetradecasaccharide in the presence of two different crowding agents, dextran, a commonly used crowding agent, as well as hyaluronan (HA), which is normally present at the cell surface. The heparin tetradecassaccaride was immobilised through its reducing end onto a biosensor surface and the crowding agents were used in solution. The results show that the observed on-rate rate of association (k_on_) decreased over two-fold with increasing concentrations of both crowding agents. The slope of initial rate of association similarly decreased, indicating that as the solution became more crowded the diffusion of FGF-2 becomes increasingly limited. One thousand kDa HA was shown to be the most effective crowding agent, as diffusion became limited at concentrations above 0.5 mg/ml, compared to 1 mg/ml and 10 mg/ml for 100kDa HA and dextran, respectively. Such concentrations of hyaluronan are readily found extracellularly. The results, therefore, suggest that the effect of crowding at the cell surface and in the extracellular matrix may play an important role in governing the interactions of at least FGF-2 with its heparan sulfate co-receptor.

## Introduction

Molecular interactions are the key to understanding the behaviour of cells. In multicellular organisms, the assembly of regulatory complexes of proteins on glycosaminoglycans in the pericellular matrix and at the cell surface represents a node of integration for the transport and effector functions of hundreds of extracellular regulatory proteins. A classic example of such interactions is that of fibroblast growth factor-2 (FGF-2) with heparan sulfate (Delehedde, *et al*. 2001). The characterisation of these regulatory interactions is required to understand molecular mechanisms and to build models of the molecular systems that regulate the growth and maintenance of multicellular organisms. The regulatory interactions are universally characterised in dilute solutions of the proteins and the glycosaminoglycan, e.g., (Delehedde, *et al*. 2002, Powell, *et al*. 2002, Rahmoune, *et al*. 1998). However, the cell surface and pericellular matrix are extremely crowded environments. Roughly in order of increasing size, extending from the cell surface is a dense network of glycolipids, glycoproteins, proteoglycans and hyaluronan (HA), making up the glycocalyx/ pericellular matrix.

Molecular crowding is considered to have two effects on molecular interactions (Ellis 2001, Minton 2001). First, non-specific interactions are greatly increased and result from transient electrostatic, steric or hydrophobic reactions between different molecules in the medium. They are weaker and of shorter duration than specific reactions, but unavoidable in physiological systems and may induce significant changes in the rate and equilibrium of reactions. Second, the excluded volume effect occurs because solute molecules are impenetrable, therefore reducing the volume of solvent available to a particular species. If the solvent volume is reduced, the effective concentration of the molecule will appear to be higher than the actual concentration when the whole volume is considered.

Crowding thermodynamically favours the binding of molecules. As molecules associate, the volume available to other species increases. The reduction in excluded volume is a more favourable state as the free energy of the solution decreases. However, it would be expected that the rate of diffusion of each molecular species would decrease due to an increased number of collisions with other molecules present. Thus if a binding reaction is diffusion limited, the rate will decrease in crowded environments (Ellis 2001, Minton 2001).

Here the binding parameters of FGF-2 for a heparin-derived tetradecasaccharide are examined in the presence of two crowding agents. Dextran is a commonly used crowding agent, since it is a long polysaccharide with some branching, and no hydrophobic patches and is regarded as inert (Ellis 2001, Minton 2001). HA must be considered as a major crowding agent of the pericellular matrix and cell surface, though it is not typically used in crowding experiments, which hitherto have focussed on intracellular proteins. HA is polyanionic, possesses considerable structure in solution and may therefore interact more with the substrates. The results demonstrate that crowding markedly reduces the on-rate of the interaction between FGF-2 and that changing the size of HA, which is a common physiological event (West 2000, West and Fan 2002), is likely to influence the diffusion of proteins in the pericellular environment.

## Results and Discussion

### Binding of FGF-2 to a heparin-derived tetradecasaccharide

The interaction between FGF-2 and a heparin tetradecasaccharide immobilised on a streptavidin-derivatised sensor surface through a reducing end biotin was chosen because it represents a model of a biologically important event and it is well characterised (Delehedde, *et al*. 2002). The binding of 20 *μ*g/ml FGF-2 to immobilized heparin-derived tetradeca-saccharide was always monophasic (result not shown), so a one site binding model was deemed to be appropriate for the analysis of the binding data. Kinetic analysis of the interaction gave values (result not shown) of the association rate constant and equilibrium dissociation rate constant similar to those obtained previously (Delehedde, *et al*. 2002) for this interaction.

### Effect of crowding agents on the binding of FGF-2

To investigate what effect molecular crowding may have on the interaction between FGF-2 and the immobilised heparin tetradecasaccharide, three binding parameters, k_on_, the extent of binding and the slope of initial rate were measured at a single concentration of FGF-2. The k_on_ or apparent on-rate depends on the association rate constant, the dissociation rate constant and the effective concentration of the ligate, FGF-2. In a surface measurement, as made in optical biosensors, k_on_ can also come to depend on the rate of diffusion of the ligate from the bulk solution into the stationary phase near the sensor surface (Fernig 2001). The extent of binding (600 arc s corresponds to 1 ng protein/mm^2^ sensor surface) is the maximum amount of FGF-2 bound at or near equilibrium and this depends on the effective concentration of FGF-2 and any competing interactions with the crowding agent. The slope of initial rate depends on the association rate constant and the effective concentration of ligate; importantly it provides an indication of whether a binding reaction is becoming diffusion limited (Edwards and Leatherbarrow 1997). Diffusion limitation at a constant ligate concentration is evidenced by a decrease in slope of initial rate.

All three crowding agents were found to affect k_on_ as the concentration of crowding agent increased (Fig. 1A). With dextran the decrease was apparent at 10 mg/mL and at 100 mg/mL dextran k_on_ was 25% of the control value. HA (100 kDa) had no effect on k_on_ until 10 mg/mL where k_on_ was 60% of the control value. Higher concentrations were not used due to the viscosity of the solutions. In contrast 1000 kDa HA was far more effective, since k_on_ was below the control value at 0. 5 mg/mL and was only 40% of the control at 5 mg/mL. Again, higher concentrations were not tested due to the viscosity of the solutions.

**Figure 1.**
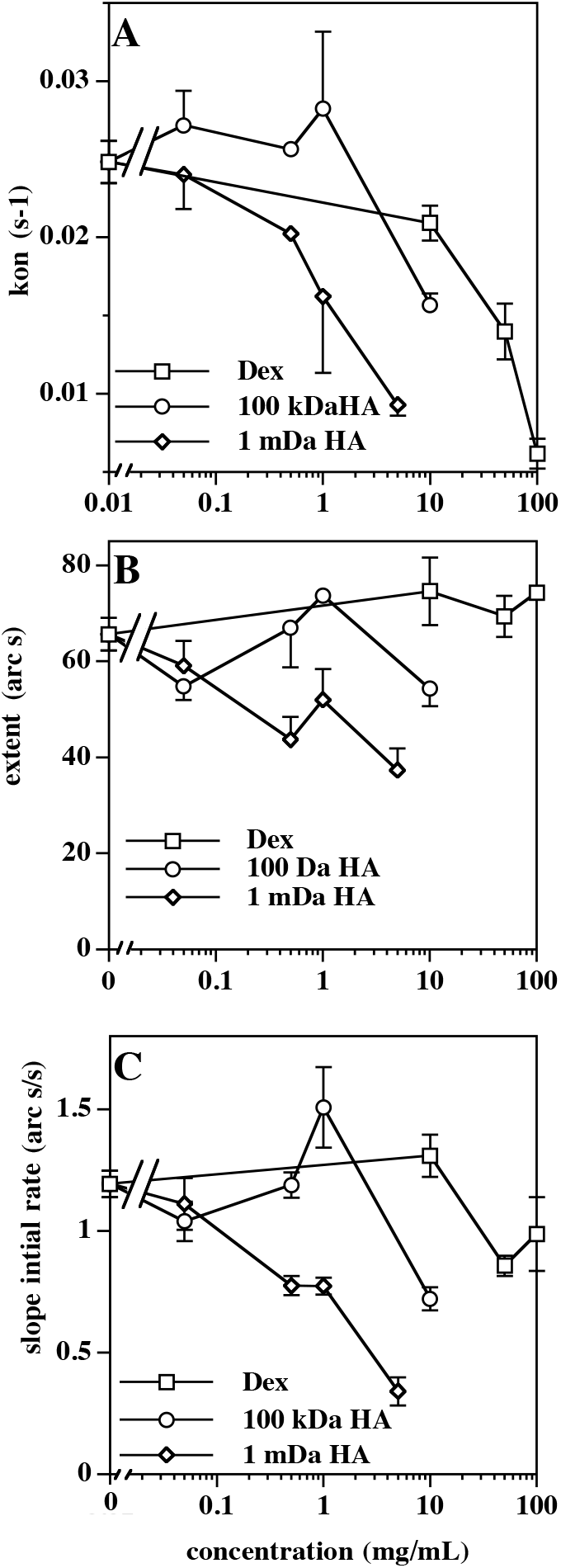
The effect of crowding agents on the binding of FGF-2 to an immobilised heparin-derived tetradecasaccharide. The k_on_, extent of binding and slope of initial rate were determined for 20 *μ*g/ml FGF-2 binding to a heparin tetradecasaccharide in the presence of various crowding agents. The three binding parameters were calculated from the binding curves as described in “Materials and Methods”. Values are the mean ± S.E. of at least three measurements except for those at 0 mg/ml, which were calculated from 25 measurements. (**A**) k_on_, (**B**) extent of binding, (**C**) slope of initial rate.

As regards the extent of binding, dextran was without effect and the small changes observed with 100 kDa HA were within the error envelope of the measurement (Fig. 1B). However, 1000 kDa HA did cause a modest (17%) but significant decrease in the extent of binding (Fig. 1B). The minimum binding structure in heparin for FGF-2 corresponds to a tetrasaccharide (Delehedde, *et al*. 2002). Given the identical chemical structures in 100 kDa HA and 1 MDa HA, it seems unlikely that the decrease in extent of binding observed with the larger HA species can be due competition resulting from a weak interaction between FGF-2 and HA, unless the 1 MDa HA adopts a solution structure in which competitive weak FGF-2 binding sites are apparent.

The slope of initial rate only decreased marginally in the presence of dextran (Fig. 1C). Intriguingly, 100 kDa HA caused a small increase in the slope of initial rate, which corresponds to the initial plateau in k_on_ (Fig. 1A) and then at 10 mg/mL the slope of initial rate fell below the control value. A clear dose-dependent decrease in slope of initial rate was observed with 1000 kDa HA (Fig. 1C).

The higher concentrations of HA are a good representation of the concentrations likely to exist in the pericellular matrix. Thermodynamically, crowding favours association reactions. At the concentrations used here, there is only weak evidence for an increase in binding reaction rate for 1 mg/ml 100 kDa HA. Instead, each crowding agent causes a decrease in k_on_. This suggests that at the concentrations used here with highly asymmetric molecules such as dextran and HA, the excluded volume effect is dominated by the vastly increased number of collisions between the FGF-2 and the crowding agent causing a decrease in diffusion rate and hence association rate. In support of this argument, evidence for a decrease in the rate of diffusion of FGF-2 is provided by the measurements of slope of initial rate, which decreases in parallel with k_on_, though not exactly synchronously.

The crowding system employed in this study is extremely simplistic compared to the pericellular matrix and the cell surface, yet the data illustrate some important points. First, the values for association rates obtained from measurements in dilute solutions may be substantially higher than the actual values for the binding events that occur in the pericellular matrix and at the cell surface and the corresponding equilibrium dissociation constant (Kd) may be markedly increased by crowding. Second, HA, which is a major species present in the pericellular matrix and at the cell surface is shown to be far more effective in reducing the rate of binding of FGF-2 to a tetradecasaccharide than dextran. This is unlikely to be simply due to molecular size, since 100 kDa HA and the dextran are of similar size yet, the 100 kDa HA has a greater crowding effect. Thus, the organisation of HA in solution may increase its effectiveness as a crowding agent. In support of this contention is the observation that 1 MDa HA is substantially more effective as a crowding agent on a weight basis than 100 kDa HA. Therefore, it is possible that modifying the size of HA in the pericellular matrix, which is well-known physiological event (West 2000, West and Fan 2002), may have dramatic effects on the binding reactions between extracellular regulatory proteins such as FGF-2 and sulphated glycosaminoglycans.

## Materials and Methods

### Materials

Aminosilane biosensor cuvettes were obtained from Thermo Electron, Basingstoke, UK. Heparin-derived tetradecasaccharide, prepared by partial heparinase I digests of pig mucosal heparin were obtained from Iduron (Manchester, UK) and biotinylated at the reducing end using biotin-XX-hydrazide (Calbiochem, Merck Biosciences Ltd, Nottingham, UK), as described (Delehedde, *et al*. 2002). Human recombinant FGF-2 was prepared as described (Ke paper). Ultrapure hyaluronan of size 100 kDa and 1000 kDa (A gift from Genzyme Advanced Biomaterials, Cambridge, USA) and T70 dextran (Sigma – Aldrich Company Ltd, Poole, UK) were dissolved in PBST (phosphate-buffered saline, 140 mM NaCl, 5 mM Na_2_HPO_4_, 5 mM NaH_2_PO_4_, pH 7.2 supplemented with 0.02 % Tween 20 (v/v).

### Binding assays

Binding reactions were carried out in aminosilane cuvettes derivatized with streptavidin and biotinylated tetradecasassharide in an IAsys two channel resonant mirror optical biosensor as described (Delehedde, *et al*. 2002, Fernig 2001, Rahmoune, *et al*. 1998). A binding assay consisted of adding 1 *μ*L FGF-2 (500 ng/ml) to a cuvette containing 24 *μ*L PBST or 24 *μ*L PBST containing the appropriate concentration of HA or dextran. The association reaction was followed for a set time, usually 180 s. The cuvette was then washed three times with 50 *μ*L PBST and the surface was then regenerated by washing twice with 50 *μ*L 2 M NaCl, 10 mM Na_2_HPO_4_, pH 7.2. Data were collected three times a second.

### Data analysis

A single binding assay yielded three binding parameters: the slope of initial rate of association, the on-rate constant (k_on_) and the extent of binding, all calculated from the association phase using the non-linear curve fitting FastFit supplied by the manufacturer (Thermo Electron). The slope of initial rate was calculated from the first 20 s of binding data, since this portion of the association reaction was linear, whereas k_on_ and the extent of binding were calculated from the entire 180 s of the association reaction. By using a relatively low level of immobilisation of biotinylated tetradecasaccharide and low concentrations of FGF-2, artefactual second phase binding sites were avoided and association data were always monophasic. Results are reported as the mean ± s.e. of three measurements, except for the values calculated for FGF-2 binding in the absence of crowding agent, which are the mean of twenty five measurements.

## Acknowledgements

The author thanks the Cancer and Polio Research Fund and the North West Cancer for financial support, David C. West for the HA and reviewing the manuscript and Claire Moran for technical assistance.

## Abbreviations

FGF-2: fibroblast growth factor-2
HA: hyaluronan.

